# Core and Matrix Thalamic Sub-Populations Relate to Spatio-Temporal Cortical Connectivity Gradients

**DOI:** 10.1101/2020.02.28.970350

**Authors:** Eli Müller, Brandon Munn, Luke J. Hearne, Jared B. Smith, Ben Fulcher, Luca Cocchi, James M. Shine

**Affiliations:** Brain and Mind Centre, The University of Sydney, Sydney, NSW, Australia; Center for Molecular and Behavioral Neuroscience, Rutgers University, Newark, USA; Molecular Neurobiology Laboratory, Salk Institute for Biological Studies, La Jolla, CA, USA; Complex Systems Research Group, The University of Sydney, Sydney, NSW, Australia; QIMR Berghofer, Brisbane, QLD 4006, Australia

## Abstract

Recent neuroimaging experiments have defined low-dimensional gradients of functional connectivity in the cerebral cortex that subserve a spectrum of capacities that span from sensation to cognition. Despite well-known anatomical connections to the cortex, the subcortical areas that support cortical functional organization have been relatively overlooked. One such structure is the thalamus, which maintains extensive anatomical and functional connections with the cerebral cortex across the cortical mantle. The thalamus has a heterogeneous cytoarchitecture, with at least two distinct cell classes that send differential projections to the cortex: granular-projecting ‘Core’ cells and supragranular-projecting ‘Matrix’ cells. Here we use high-resolution 7T resting-state fMRI data and the relative amount of two calcium-binding proteins, parvalbumin and calbindin, to infer the relative distribution of these two cell-types (Core and Matrix, respectively) in the thalamus. First, we demonstrate that thalamocortical connectivity recapitulates large-scale, low-dimensional connectivity gradients within the cerebral cortex. Next, we show that diffusely-projecting Matrix regions preferentially correlate with cortical regions with longer intrinsic fMRI timescales. We then show that the Core–Matrix architecture of the thalamus is important for understanding network topology in a manner that supports dynamic integration of signals distributed across the brain. Finally, we replicate our main results in a distinct 3T resting-state fMRI dataset. Linking molecular and functional neuroimaging data, our findings highlight the importance of the thalamic organization for understanding low-dimensional gradients of cortical connectivity.

## Introduction

Advances in whole-brain neuroimaging have identified low-dimensional gradients in the cerebral cortex that interconnect sensorimotor regions with limbic structures (Margulies et al., 2016; Mesulam, 1998). Recent work has shown that these same gradients are underpinned by structural differences that have long been known to comprise the cerebral cortex (García-Cabezas et al., 2019), including circuit complexity (Paquola et al., 2019; Vázquez-Rodríguez et al., 2019) and the extent of cortical myelination (Burt et al., 2018; Demirtaş et al., 2019). Circuit complexity and myelination are presumed to differentially enforce coupling between structure and function (Fallon et al., 2020), with granular sensory regions more closely tethered to their structural constraints than associative agranular regions (Paquola et al., 2019; Vázquez-Rodríguez et al., 2019). The complexity of cortical circuitry has also been shown to underpin a gradient of temporal scales across the brain, with associative regions fluctuating across relatively longer time scales than sensory regions (Honey et al., 2012). Comparable low-dimensional gradients have also been observed in task contexts (Shine et al., 2019a) and share similarities with meta-analytic studies of task fMRI (Margulies and Smallwood, 2017), suggesting that the low-dimensional gradients in the cerebral cortex may be a crucial factor underlying whole-brain functional organization.

A phylogenetic perspective on the brain (Cisek, 2019) suggests that the cortex is supported by several subcortical structures that shape, constrain, and augment its activity on a moment-to-moment basis. One such structure that is crucial for shaping whole brain dynamics is the thalamus (Halassa and Sherman, 2019; Jones, 2009; 2001; Shine et al., 2019b) (Figure 1). Located in the diencephalon, the thalamus is reciprocally connected with the entire cerebral cortex, along with primary sensory receptors (such as the retina and dorsal column tract) and numerous other subcortical systems (such as the basal ganglia, superior colliculus and cerebellum). These inputs innervate distinct, anatomically-segregated sub-nuclei within the thalamus (Figure 1) that are surrounded by the shell-like GABAergic reticular nucleus (Jones, 2001). Through activity-dependent GABAergic inhibition of cortically- and subcortically-driven activity, the thalamus likely plays a crucial role in shaping and constraining patterns of whole-brain dynamics (Halassa and Sherman, 2019; Jones, 2009; 2001).

**Figure 1.**
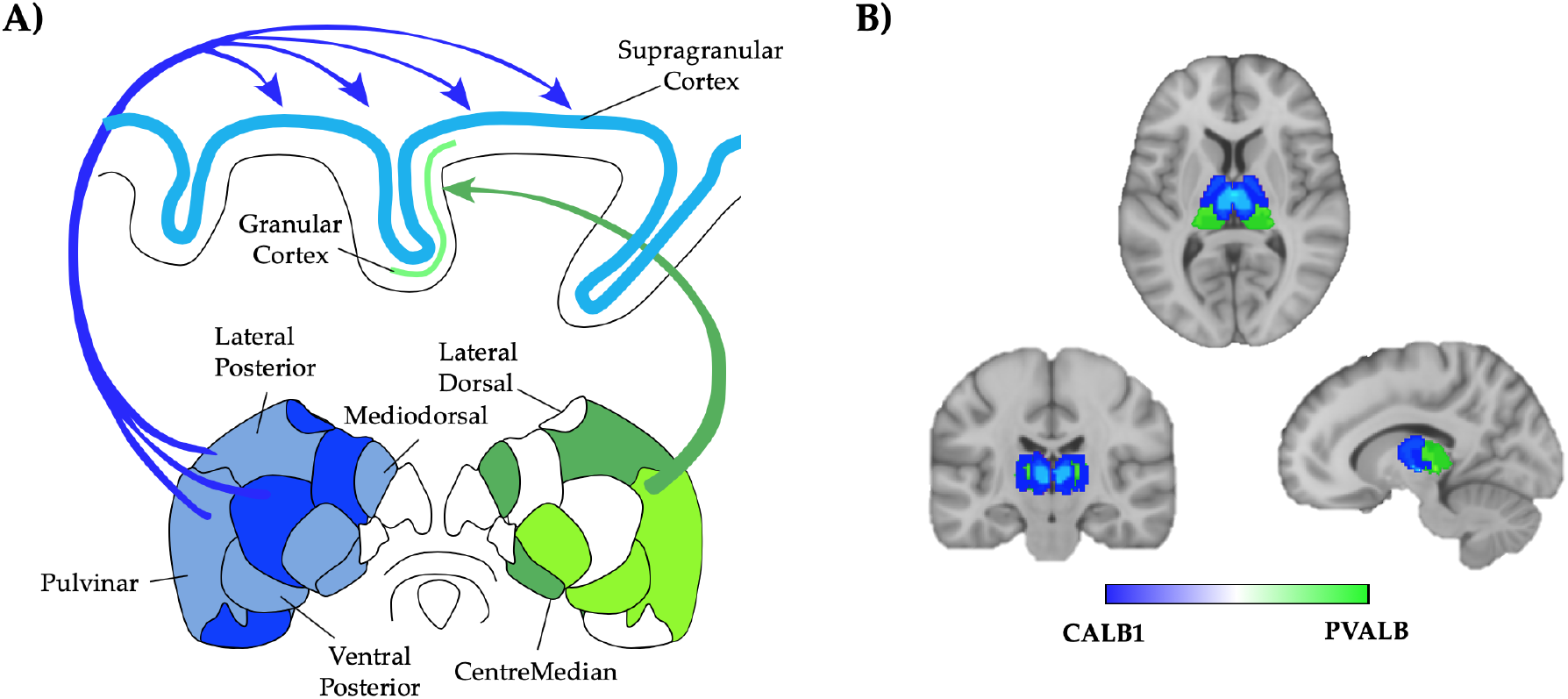
The Cytoarchitecture of the Thalamus. A) Matrix (blue) and Core (green) thalamic populations have distinct projection patterns: Core cells innervate granular layers of the cerebral cortex in a specific fashion, whereas Matrix cells innervate supragranular cortex in a relatively diffuse manner. Color intensity denotes the relative expression of CALB1 and PVALB in different thalamic subnuclei (Jones, 2009; 2001); B) Axial (upper), coronal (lower left) and sagittal (lower right) loading of calbindin values (CALB1; blue) and parvalbumin values (PVALB; green) within the thalamus.

Individual sub-nuclei within the thalamus innervate distinct regions of the cerebral cortex (Figure 1A) (Jones, 2009; 2001). In addition, each thalamic sub-nucleus is comprised of a mixture of excitatory neurons that project in unique ways to the cerebral cortex (Clascá et al., 2012; Jones, 2009). Specifically, Parvalbumin-rich ‘Core’ cells (Figure 1B; green), typically dense within sensory thalamic nuclei such as the lateral geniculate nucleus, send axonal projections to Layers III and IV of the cerebral cortex. These cells are thought to act as ‘drivers’ of feed-forward activity (Jones, 2009; 2001). In contrast, Calbindin-rich ‘Matrix’ cells (Figure 1; blue), which are more prevalent in higher-order thalamic nuclei (Herkenham, 1979; Jones, 2001), preferentially target agranular cortices in a more distributed fashion that crosses typical cortical receptive fields (Jones, 2009; 2001). Whereas individual thalamic nuclei contain a blend of both cell-types, some nuclei are almost exclusively of the ‘Matrix’ type (these are known as ‘intralaminar’ nuclei; (Van der Werf et al., 2002). Of note, other schemes have been devised to parse the thalamus (Phillips et al., 2019), including the class of cortical region proposed to ‘drive’ activity in the glutamatergic relay nuclei (Sherman, 2007). However, these proposals are not considered in detail here.

The densities of Parvalbumin- (Core) and Calbindin- (Matrix) neurons have been used to classify the relative proportion of cells within each thalamic nucleus that project to either granular or supragranular layers, respectively (Figure 1A) (Jones, 2009; 2001; Münkle et al., 2000). These studies have provided anatomical evidence for the presence of an inter-digitated cellular architecture within the thalamus. The presence of calcium-binding proteins has also been shown to delineate further levels of anatomical specificity in the thalamus, such as the extent and location of laminar projections to the cortex (Clascá et al., 2012). However, the importance of this anatomical organizing principle for shaping patterns of whole-brain activity and connectivity has been less well characterized.

In this work, we leveraged the high spatiotemporal resolution of a 60-subject, 7T resting-state fMRI dataset to extract time series from the thalamus to examine the relationship between thalamic cell-population densities and whole-brain functional connectivity. We first calculated the time-averaged connectivity between each thalamic voxel and 400 cortical parcels from the Schaefer atlas (Schaefer et al., 2018). We then used the Allen Human Brain Atlas (Gryglewski et al., 2018; Hawrylycz et al., 2012) to estimate the expression of two calcium-binding proteins – PVALB and CALB1 – in each thalamic voxel. This allowed us to determine the relative weighting of Core versus Matrix cells, respectively (Jones, 2009; 2001; Münkle et al., 2000; Phillips et al., 2019). These thalamic variations were then compared to fluctuations in the activity of parcels within the cerebral cortex, allowing us to quantify the relationship between patterns of thalamic activity and emergent patterns of whole-brain functional connectivity.

## Methods

### Participants

Sixty-five healthy, right-handed adult participants (18–33 years) were recruited, of whom 60 were included in the final analysis (28 females). Participants provided informed written consent to participate in the study. The research was approved by The University of Queensland Human Research Ethics Committee. These data were originally described in Hearne et al., 2017.

### Neuroimaging Acquisition

1050 (~10 minutes) whole-brain 7T resting state fMRI echo planar images were acquired using a multiband sequence (acceleration factor = 5; 2 mm^3^ voxels; 586 ms TR; 23 ms TE; 40^0^ flip angle; 208 mm FOV; 55 slices). Structural images were also collected to assist functional data pre-processing (MP2RAGE sequence – 0.75 mm^3^ voxels 4,300 ms TR; 3.44 ms TE; 256 slices).

### Data Pre-processing

DICOM images were first converted to NIfTI format and realigned. T1 images were reoriented, skull-stripped (FSL BET), and co-registered to the NIfTI functional images using statistical parametric mapping functions. Segmentation and the DARTEL algorithm were used to improve the estimation of non-neural signal in subject space and the spatial normalization. From each gray-matter voxel, the following signals were regressed: linear trends, signals from the six head-motion parameters (three translation, three rotation) and their temporal derivatives, white matter, and CSF (estimated from single-subject masks of white matter and CSF). The aCompCor method (Behzadi et al., 2007) was used to regress out residual signal unrelated to neural activity (i.e., five principal components derived from noise regions-of-interest in which the time series data were unlikely to be modulated by neural activity). Participants with head displacement > 3 mm in > 5% of volumes in any one scan were excluded (*n* = 5). A temporal band pass filter (0.071 < *f* < 0.125 Hz) was applied to the data. Following pre-processing, the mean time series was extracted from 400 predefined cortical parcels using the Schaefer atlas (Schaefer et al., 2018).

### Relative Expression of Calbindin and Parvalbumin in the Thalamus

We co-registered the thalamic Morel atlas (Niemann et al., 2000) to MNI152 space and then identified 2305 voxels across both hemispheres that were inclusive to the thalamic mask. We then used this volume as a mask to extract the mRNA level provided by the Allen Human Brain Atlas (Hawrylycz et al., 2012). Prior to extraction, data were smoothed using variogram modelling (Gryglewski et al., 2018). We extracted spatial maps of estimated mRNA levels for two genes that are known to express distinct calcium-binding proteins: Calbindin (CALB1) and Parvalbumin (PVALB). These proteins have been previously shown to delineate two distinct subpopulations of thalamic projection cells (Matrix and Core, respectively) with distinct anatomical connectivity profiles. Note that there are other calcium binding proteins (notably, Calretinin) with substantial expression in the thalamus (Münkle et al., 2000; Phillips et al., 2019), however these patterns are not considered here.

To create a realistic estimate of the relative weighting of each thalamic sub-population, these values were first normalized (z-score) across all voxels within the thalamic mask before creating the difference between the normalized Calbindin and Parvalbumin values (Figure 1B). We denote this measure CP_*T*_: positive values reflect voxels with higher CALB1 values, negative values reflect voxels with higher PVALB values and values of zero denote a balance between the two populations. To further validate the measure, the mean CP_*T*_ value was calculated for each of 31 pre-defined thalamic subnuclei (Table 1) and compared qualitatively to results obtained from direct histological (Münkle et al., 2000) and immunohistochemistry (Arai et al., 1994) analyses. We observed a positive correlation between these direct measurements and our geneexpression based proxy, CP_*T*_ (*r* = 0.550; *p* = 0.003; Figure S1).

**Table 1.** Thalamic Morel Atlas and Calcium Binding Protein Values. mRNA loading of Calbindin (CALB1) and Parvalbumin (PVALB) for each of 31 pre-identified thalamic sub-nuclei, along with the X, Y and Z MNI Coordinates for the center of each region.

| Nucleus | CALB1 | PVALB | X | Y | Z |
| --- | --- | --- | --- | --- | --- |
| Anterior Ventral | 2.0 | 1.8 | 61.2 | 45.8 | 42.6 |
| Lateral Dorsal | 8.3 | 6.2 | 55.2 | 46.3 | 44.8 |
| Magnocellular Lateral Geniculate | 6.5 | 5.2 | 50.9 | 48.6 | 33.3 |
| Parvocellular Lateral Geniculate | 2.3 | 1.9 | 51.7 | 46.5 | 34.0 |
| Medial Geniculate | 6.0 | 3.7 | 50.4 | 45.4 | 34.6 |
| Anterior Ventral Posterior Lateral | 12.2 | 7.2 | 54.1 | 44.5 | 39.4 |
| Posterior Ventral Posterior Lateral | 11.8 | 7.3 | 53.5 | 45.7 | 41.0 |
| Medial Ventral Posterior Lateral | 9.7 | 6.5 | 54.0 | 45.5 | 36.9 |
| Inferior Ventral Posterior Lateral | 9.7 | 5.6 | 52.3 | 45.7 | 36.2 |
| Anterior Ventral Lateral | 9.0 | 7.2 | 58.6 | 45.9 | 39.5 |
| Dorsal Posterior Ventral Lateral | 9.4 | 7.4 | 57.8 | 45.7 | 43.9 |
| Ventral Posterior Ventral Lateral | 10.1 | 7.6 | 57.0 | 45.4 | 39.6 |
| Magnocellular Ventral Anterior | 1.5 | 3.1 | 61.1 | 46.4 | 37.9 |
| Magnocellular Ventral Posterior | 2.7 | 3.1 | 61.4 | 45.6 | 39.9 |
| Ventral Medial | 6.9 | 7.5 | 57.7 | 45.7 | 36.4 |
| Magnocellular Mediodorsal | 5.4 | 6.3 | 57.1 | 45.6 | 39.1 |
| Parvocellular Mediodorsal | 8.1 | 7.5 | 56.2 | 45.6 | 40.1 |
| Medial Pulvinar | 9.2 | 5.8 | 49.1 | 45.7 | 39.4 |
| Lateral Pulvinar | 3.5 | 2.0 | 49.4 | 46.3 | 38.7 |
| Anterior Pulvinar | 12.0 | 7.4 | 52.2 | 46.9 | 39.3 |
| Limitans | 8.7 | 6.4 | 50.8 | 45.8 | 37.0 |
| Lateral Posterior | 10.4 | 7.0 | 51.9 | 44.1 | 43.5 |
| Central Lateral | 9.1 | 7.2 | 54.9 | 45.9 | 41.4 |
| CentroMedian | 2.5 | 3.5 | 59.4 | 45.4 | 37.1 |
| Central Median | 9.8 | 7.3 | 53.7 | 45.4 | 37.6 |
| Parafascicular | 6.2 | 5.2 | 54.3 | 45.4 | 36.1 |

### Thalamo-Cortical Connectivity Patterns

Following the creation of the CP_*T*_ spatial map, Pearson’s correlations were calculated between each thalamic voxel and the mean timeseries of the 400 cortical parcels for all 60 subjects. For each cortical parcel, we then correlated its functional connectivity to each voxel within the thalamic mask with CP_*T*_. This allowed us to quantify, with a single value (CP_*C*_), the relative connectivity between each of the 400 cortical parcels and either the Matrix or Core thalamic population activity. As CP_*T*_ represents the relative expression of CALB1 (Matrix-rich) versus PVALB (Core-rich), positive correlations (i.e., CP_*C*_ > 0) were interpreted as preferential functional coupling with Matrix thalamic populations and negative correlations (i.e., CP_*C*_ < 0) were taken to be indicative of preferential coupling to Core populations. To aid interpretation, CP_*T*_ was projected onto a thalamic volume (Figure 1), CP_*C*_ was projected onto the cortical surface (Figure 2), and the mean correlation value within each of 17 pre-defined resting-state networks was calculated (Schaefer et al., 2018).

**Figure 2.**
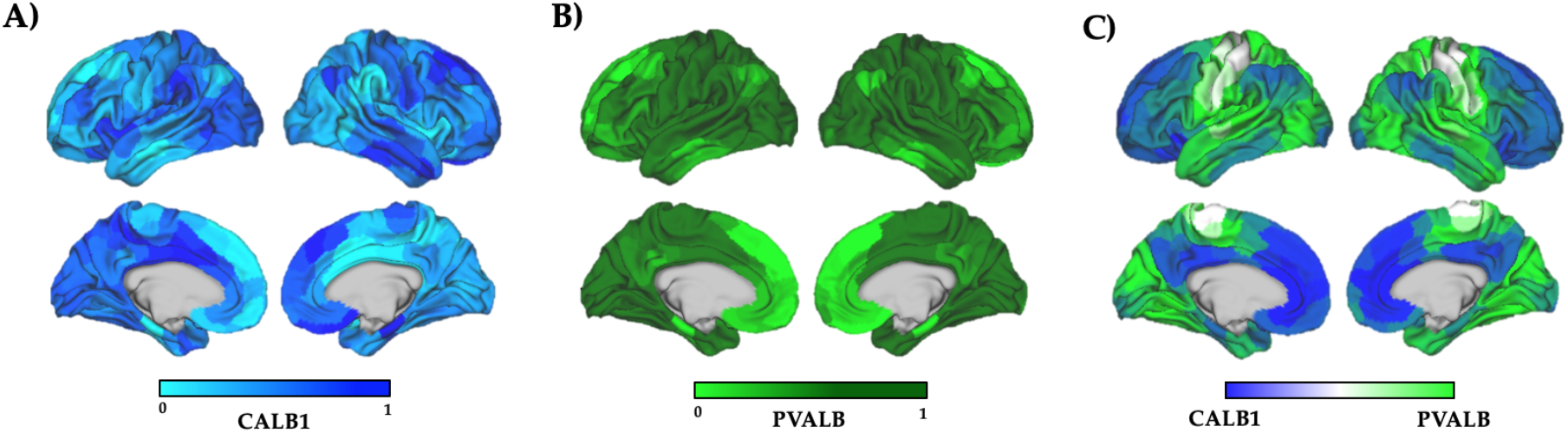
Correlations between Matrix-Core thalamic regions and cortical parcels, CP_*C*_. A) standardized correlation between Matrix thalamus (CALB1 levels) and cortical parcels; B) standardized correlation between Core thalamus (PVALB levels) and cortical parcels; C) relative difference between Matrix and Core populations relationship with cortical parcels. Colour bars depict the relative strength of correlation between each thalamic population at the individual cortical parcels (standardized between 0 and 1).

### Relationship with Low-Dimensional Cortical Gradients

Given the known diverse connectivity of the thalamus with the cerebral cortex, we predicted that known anatomical gradients in the thalamus should relate to lowdimensional patterns of functional connectivity in the cortex. To test this hypothesis, we compared the Core–Matrix organization with the low-dimensional cortical gradients obtained by applying diffusion embedding analyses to resting-state fMRI data from a large cohort of subjects (Margulies et al., 2016). Un-thresholded volumetric maps representing the top 5 low-dimensional cortical gradients were downloaded from NeuroVault (Gorgolewski et al., 2015). These maps were then down sampled into the 400 cortical parcel space by calculating the mean expression of each gradient within each cortical parcel. We then compared the CP_*C*_ connectivity pattern to each of the top five diffusion embedding components using Pearson’s correlations (statistical testing utilized parametric, one-sided ‘spin’ tests to control for nonlinearity and spatial autocorrelation, respectively; see below for details).

### Relationship Between CP_*C*_ and Intrinsic Timescale

Different areas of the cerebral cortex are characterized by an intrinsic timescale that reflects the length of the time window over which the signals entering each brain region are integrated (Mesulam, 1998; Honey et al., 2012). There are two predominant methods utilised in fMRI studies for investigating the intrinsic timescale of activity: autocorrelation (Murray et al., 2014; Watanabe et al., 2019) and fractal based (He, 2011; Churchill et al., 2016) techniques. While both measures identify temporal selfsimilarity, autocorrelation techniques typically focus on linear correlations with past values of a particular region, whereas in contrast, fractal analysis is used as a measure of complexity (Dong et al., 2018) and can be used to discern the significance of differing time-scale frequencies. For instance, a purely fractal signal suggests all frequencies contain information (Churchill et al., 2016).

To examine the relationship between Core–Matrix architecture and the intrinsic BOLD timescale of cortical parcels, we first performed an autocorrelation analysis (Murray et al., 2014). Specifically, to estimate the extent of the autocorrelation, *ac*, in each region, *r*, we calculate the correlation coefficient between the BOLD activity, *X*_r_ (*t*), at times *t* and *t* + *k* with different time lags *k* = 0, 1, …, *k*_max_. For our analysis *k*_max_ = 50 TR, however our findings were not sensitive for differences in *k*max (from 15 – 1000). The autocorrelation for each region is given by

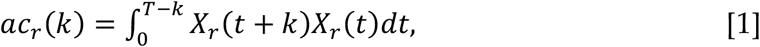

where *T* is the duration of the BOLD activity and the autocorrelation is normalised such that the autocorrelation at a lag of zero is unity. As the time lag increases, the autocorrelation decays according to the regions’ intrinsic timescale, which is well fit to an exponential decay, *N_r_* with an offset given by

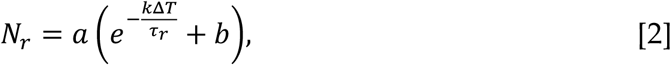

where *τ_r_* is the intrinsic timescale, *a* and *b* are the scaling and offset constants, respectively, *k* is the lag, and Δ*T* is the time resolution (i.e., the length of the TR). We fit the exponential decay at a population level, where the autocorrelation is averaged across time, in order to obtain a population level statistic for each region in our dataset. The curve parameters were estimated using a Levenberg-Marquardt nonlinear leastsquares fitting algorithm. It should be noted that our method is identical to the method adopted by (Murray et al., 2014), who also fit an exponential decay with a baseline constant. Another proportional method is to calculate the area under the normalized autocorrelation curve (Gollo, 2019; Watanabe et al., 2019). In practice, we found our method was more flexible for fitting rapidly decaying BOLD signals.

To further relate the thalamocortical properties with resulting temporal dynamics, we calculated the fractal nature of the timeseries of each cortical region across all subjects. To do so, we employed a detrended fluctuation analysis (DFA) to BOLD activity at each region (Peng et al., 1995). DFA is an efficient fractal measure to estimate the Hurst exponent of a signal that is intrinsically robust to signal non-stationarity, which may be present in BOLD activity. To calculate the DFA, we first computed the integrated time-series 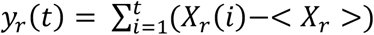, where < *X_r_* > is the average BOLD activity. We then subdivided *y_r_*(*t*) into windows of equal length *n* and estimated a linear regression per subdivision, *w_n_* (*t*). We then computed the root-mean-square magnitude of fluctuations, *F*(*n*), on the detrended data:

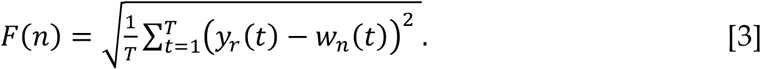

Finally, we calculated the RMS fluctuation for varying window sizes and calculated the slope of *F*(*n*) vs *n* plotted on a logarithmic scale, again using a Levenberg-Marquardt nonlinear least-squares fitting algorithm. The slope of log-log plot corresponds to the Hurst exponent, *H*, where *H* = 0.5 corresponds to Brownian motion (Beran, 1992; Munn et al., 2020). As *H* increases, the level of positive long-range temporal correlation in the signal increases. That is, an increase in BOLD activity is typically followed by another increase, and a reduction is typically followed by further reduction. As such, larger values of *H* indicate longer time scales of activity within individual regions. In our analysis, we calculated the Hurst exponent over window ranges from 5-60 TRs, sampled in logarithmically uniformly window sizes. The window range avoids zero residuals across the shortest intervals and low-frequency confounds identified in the power-spectral density across the long intervals. Consistent with (Churchill et al., 2016; He, 2011), we observed a crossover between two fractal ranges, consisting of an initial large increase of *F*(*n*) and large *H* for small window lengths due to serial autocorrelations which decay rapidly transitioning to smaller *H* values.

### Network Topology

To embed the CP_*C*_ results within the context of the broader cortical network, we estimated time-averaged functional connectivity in all 60 subjects and then subjected the weighted, un-thresholded connectivity matrices to topological analysis. This involved using a weighted- and signed-version of the Louvain modularity algorithm from the Brain Connectivity Toolbox (Rubinov and Sporns, 2010). The Louvain algorithm iteratively maximizes the modularity statistic, *Q*, for different community assignments until the maximum possible score of *Q* has been obtained (Equation 4). The modularity estimate for a given network is, therefore, quantification of the extent to which the network may be subdivided into communities with stronger within-module than between-module connections.

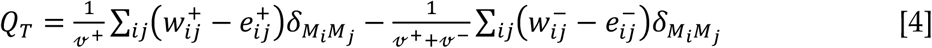

where *v* is the total weight of the network (sum of all negative and positive connections), *w_ij_* is the weighted and signed connection between regions *i* and *j*, *e_ij_* is the strength of a connection divided by the total weight of the network, and *δ_MiMj_* is set to 1 when regions are in the same community and 0 otherwise. ‘+’ and ‘–’ superscripts denote all positive and negative connections, respectively. In our experiment, the γ parameter was set to 1.1 (tested within a range of 0.5–2.0 for consistency across 100 iterations).

The participation coefficient quantifies the extent to which a region connects across all modules. This measure has previously been used to characterize diversely connected hub regions within cortical brain networks (e.g., see Power et al., 2013). Here, the Participation Coefficient (B_*iT*_) was calculated for each of the 400 cortical parcels for each subject, where κ_isT_ is the strength of the positive connections of region *i* to regions in module *s*, and κ_iT_ is the sum of strengths of all positive connections of region *i*. The participation coefficient of a region is therefore close to 1 if its connections are uniformly distributed among all the modules and 0 if all of its links are within its own module:

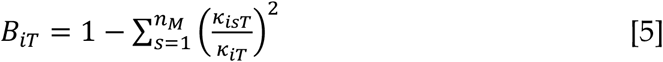

To investigate the topological relationships between the Core and Matrix populations and network topology, we classified cortical regions into three distinct groups: Core (CP_*C*_ < –0.1), Matrix (CP_*C*_ > 0.1) and participatory (parcels with participation greater than 0.5). We observed no overlap between these three groups and tested the significance of this null relationship using a permutation test (5,000 permutations, p < 0.001). For interpretation, we also plotted these three populations onto a force-directed graph (Figure 5).

### Time-varying Topological Dynamics

To quantify the relationship between CP_*C*_ and time-varying network topology, we used a sliding-window Pearson’s correlation design to estimate windowed patterns of undirected functional connectivity between all 400 cerebral cortical. In contrast to typical sliding-window approaches (Lurie et al., 2020), we split regional time series data into 20 non-overlapping windows (each of 50 TRs). This approach was adopted so as to avoid artificial smoothing that occurs with strongly overlapping windows. After averaging across multiple jitters of this tensor (+/- 5-20 TRs), we next calculated the mean of the temporal standard deviation of the results 3d tensor. The resultant vector allowed us to quantify the variability of functional connectivity measures at the regional level. We then correlated this measure with the CP_*C*_ vector to test the hypothesis that Matrix-supported cortical regions demonstrated a greater diversity of connectivity patterns than Core-supported cortical regions.

### Statistical Approach

We used non-parametric testing to determine statistical significance of the relationships identified across our study (Nichols and Holmes, 2002). A distribution of 5,000 Pearson’s correlations was calculated for each comparison, against which the original correlation was compared. Using this approach, p_*RAW*_ was calculated as the proportion of the null distribution that was less extreme than the original correlation value.

We also conducted a separate set of statistical tests in an attempt to control for the prominent spatial autocorrelation associated with low-dimensional gradients in the brain. Specifically, for each correlation, the gradients were projected back into FreeSurfer space, and the cortical parcels were rotated around the sphere (i.e., preserving spatial autocorrelation) by creating a random starting point (i.e., a value between 1 – 32,492) and re-orienting the data to this location (Reardon et al., 2018). The mean value of each of the randomly rotated values was then calculated for each parcel and used to estimate a null distribution of correlations. A distribution of 1,000 Pearson’s correlations was calculated for each comparison, against which the original correlation was compared. This statistical test was denoted as p_*SPIN*_, and was calculated as the proportion of the null distribution that was less extreme than the original correlation value.

### Reproducibility

To test the reproducibility of our results, we performed a separate replication analysis on 3T resting state fMRI data from the Human Connectome Project (100 subjects). Information regarding the specific imaging parameters from this data is presented elsewhere (Glasser et al., 2013). Data were pre-processed in the same fashion as the above analysis. Due to the fact that the comparisons were purely for replication purposes, statistical analyses were limited to traditional permutation tests (i.e., p_*RAW*_).

## Results

### Thalamic Sub-populations are Differentially Correlated with the Cerebral Cortex

We observed differential functional coupling between thalamic populations, dissociated based on gene expression, and the cortex, CP_*C*_ (Figure 2). Matrix regions of thalamus, which have higher expression of CALB1 (Figure 1B), showed a strong preference for connectivity within trans-modal cortices, incorporating Default, Control, Limbic, and Ventral Attention sub-networks (Fig. 2A, blue). In contrast, the PVALB-rich Core cells showed preferential coupling to primary somatosensory cortices, as evidenced by their stronger correlations with parcels from the Visual, Somatomotor, Dorsal-Attention and Temporo-Parietal networks (Figure 2B, green). Figure 2C highlights the difference between Matrix (blue) and Core (green) connectivity patterns.

### Matrix-Core Thalamus Recapitulates Primary Cortical Gradient

We observed a strong positive correlation between the first diffusion embedding gradient estimated from resting state fMRI data (Margulies and Smallwood, 2017) and the extent with which cortical parcels were preferentially connected with Matrix vs. Core thalamic voxels (r = 0.772, p_*RAW*_ = 2.48×10^-80^, p_*SPIN*_ = 1.0×10^-3^; Figure 3; see Figure S2 for higher gradients). Consistent with these patterns, we observed a moderate negative correlation with t1w:Tt2w ratio (r = –0.319; p_*RAW*_ = 6.6×10^-11^, p_*SPIN*_ = 0.025; Figure 3B), which is an indirect proxy for cortical myelination (Glasser et al., 2016) that is known to be negatively correlated with the hierarchical level of the cerebral cortex (Burt et al., 2018).

**Figure 3.**
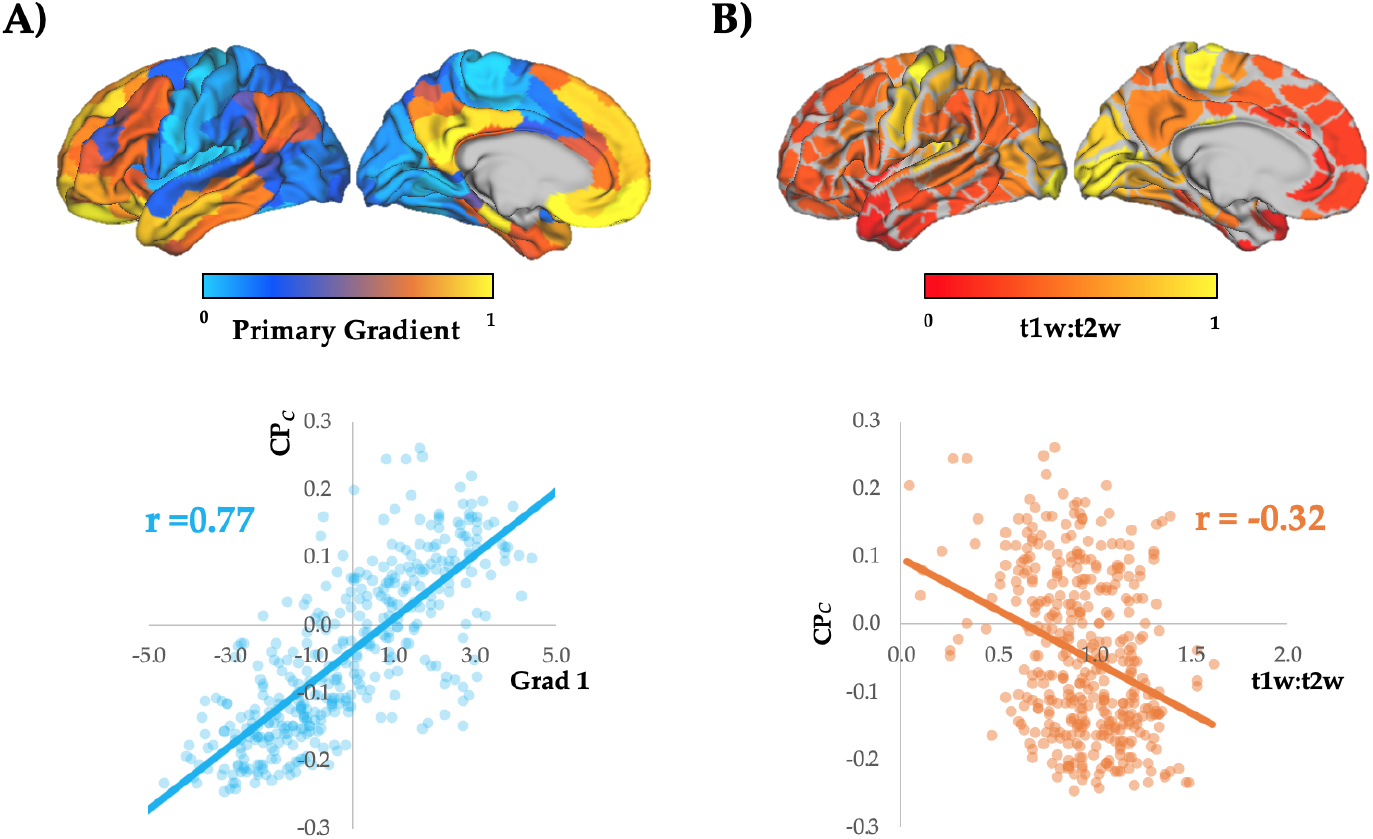
Relationship between CP_*C*_ and low-dimensional cortical gradients. A) Surface plot of the first low-dimensional gradient (Margulies and Smallwood, 2017). We observed a positive Pearson’s correlation with the CP_*C*_ (r = 0.772; p = 2.48×10^-80^); B) Surface plot of the t1w:t2w ratio on the cerebral cortex. We observed an inverse correlation with the CP_*C*_ (r = −0.320; p = 5.8×10^-11^).

### Matrix Thalamus Preferentially Couples to Regions with Slower Timescales

We observed a significant positive correlation between CP_*C*_ and both: the intrinsic timescale, *τ* (mean = 3.52±0.3, r = 0.419, p_*RAW*_ = 1.9 x 10^-18^; p_*SPIN*_ = 1.0×10^-3^; Fig 4A/B) and regional fractality, *H* (mean = 0.78±0.03, r = 0.245, p_*RAW*_ = 7.2×10^-7^, p_*SPIN*_ = 0.045; Fig 4C/D). Intuitively, we found that regional *H* and *τ* values were strongly positively correlated (r = 0.622, p_*RAW*_ = 4.0 x 10^-44^, p_*SPIN*_ = 1.0×10^-3^), suggesting that regions the longer integration windows in associative regions of the cerebral cortex (Honey et al., 2012) may be due to an increase in regional fractality (i.e., larger *H* value; Dong et al., 2018) and the presence of long-range correlations. Both timescale measures were also positively correlated with the first gradient of connectivity present within the cerebral cortex (*τ*: r = 0.598, p_*RAW*_ = 3.61 x 10^-40^, p_*SPIN*_ = 1.0 x 10^-3^; *H*: r = 0.272, p_*RAW*_ = 3×10^-8^; p_*SPIN*_ = 0.005).

**Figure 4.**
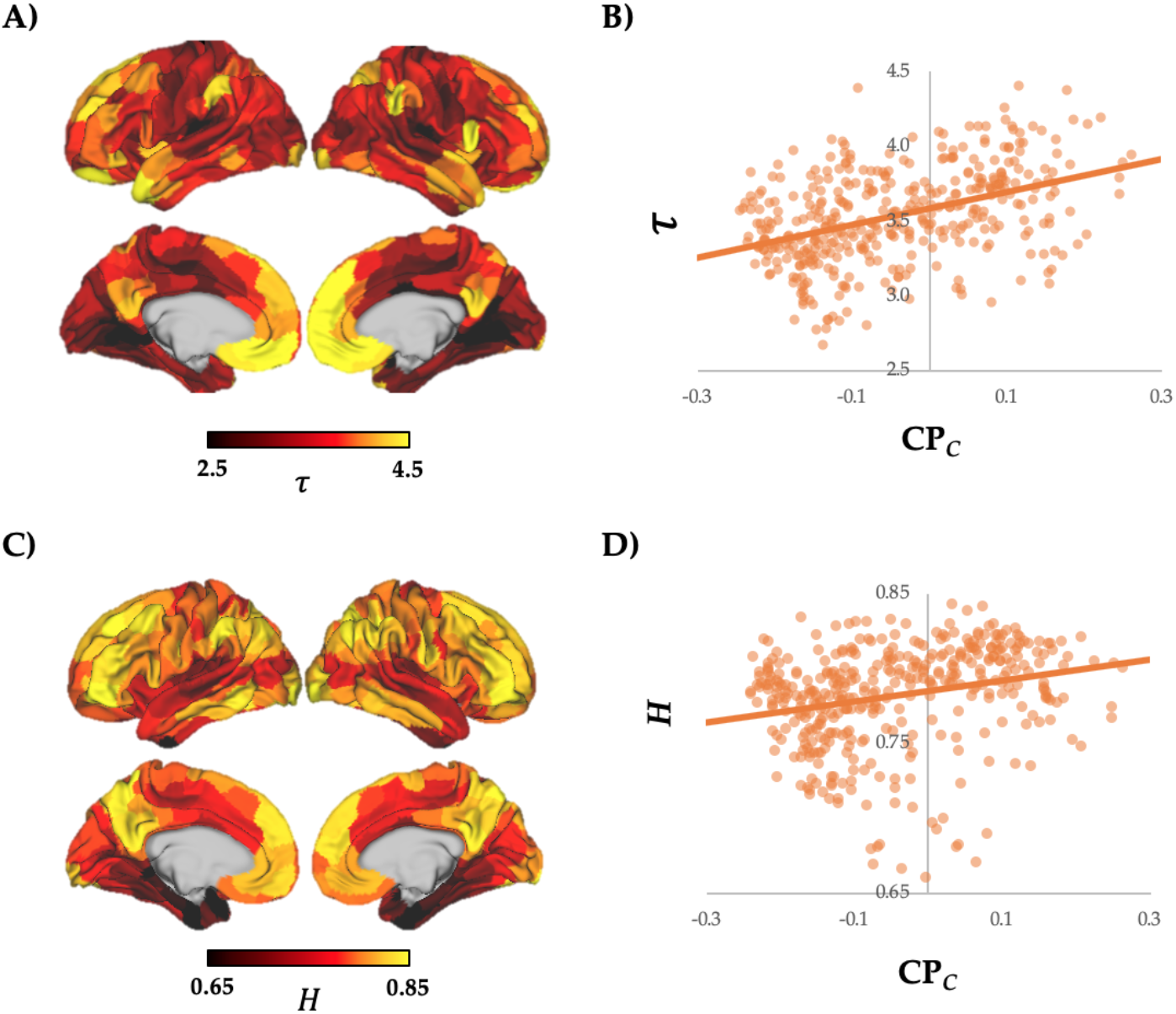
Relationship between Thalamic Connectivity and Cortical Intrinsic Timescale. A) the autocorrelation parameter, *τ*, plotted onto the cerebral cortex; B) a scatter plot between *τ* and CP_*C*_ (*r* = 0.419, p_*RAW*_ = 1.9 x 10^-18^; p_*SPIN*_ = 1.0×10^-3^); C) the Hurst exponent, *H*, plotted onto the cerebral cortex; D) a scatter plot between *τ* and CP_*C*_ (*r* = 0.245, p*_RAW_* = 7.2 x 10^-7^; p_*SPIN*_ = 0.045).

Together, these results suggest that the associative cortical regions preferentially supplied by the Matrix thalamus were associated with a longer intrinsic timescale when compared to the regions supported by the Core thalamus, at least during the relatively quiescent resting state, and may support the notion of quasi-criticality in the resting brain (Moretti and Munoz, 2013). Consistent with this hypothesis, we observed an inverse correlation between CP_*C*_ and regional Lempel-Ziv complexity (mean = 99.24±0.5, r = −0.407, p_*RAW*_ = 1.9 x 10^-17^, p_*SPIN*_ = 0.013). To test whether this relationship may be related to the complexity of thalamic timeseries, we related CP_*T*_ (i.e., CALB1:PVALB ratio in thalamic nuclei) to the Hurst exponent calculated on thalamic timeseries. The resultant correlation was positive (r = 0.366, p = 0.042), suggesting that Matrix thalamic nuclei were active over longer timescales, and hence, were responsible for promoting longer-timescale interactions in associative regions of the cerebral cortex.

### Relationship between Core and Matrix Thalamic Populations and Network Topology

Based on the known importance of the thalamus for defining whole-brain network topological integration (Bell and Shine, 2015; Hwang et al., 2017), we next compared CP_*C*_ values with measures that quantify the topology of the resting brain. We did not observe a significant correlation between CP_*C*_ and B_*T*_ (r = −0.022; p_*RAW*_ = 0.663; p_*SPIN*_ = 0.603), and although there was a weak inverse correlation with W*T* (r = −0.164, p_*RAW*_ = 0.001, p_*SPIN*_ = 0.095), it did not survive correction for spatial autocorrelation (i.e, p_*SPIN*_ > 0.05). Despite the lack of strong linear relationships between CP_*C*_ and network topology, viewing the data within a force-directed plot highlighted a key topological relationship between CP_*C*_ and B_*T*_ (Figure 5A). Namely, highly integrative regions (B_*T*_ > 50^th^ percentile; Figure 5A; red) were found to be inter-digitated between the populations of cortical parcels that were preferentially connected to either Matrix (Figure 5A; blue) or Core (Figure 5A; green) parcels. This suggests that highly integrative regions of the cerebral cortex topologically inter-connect regions on either end of the Core-Matrix thalamocortical gradient.

**Figure 5.**
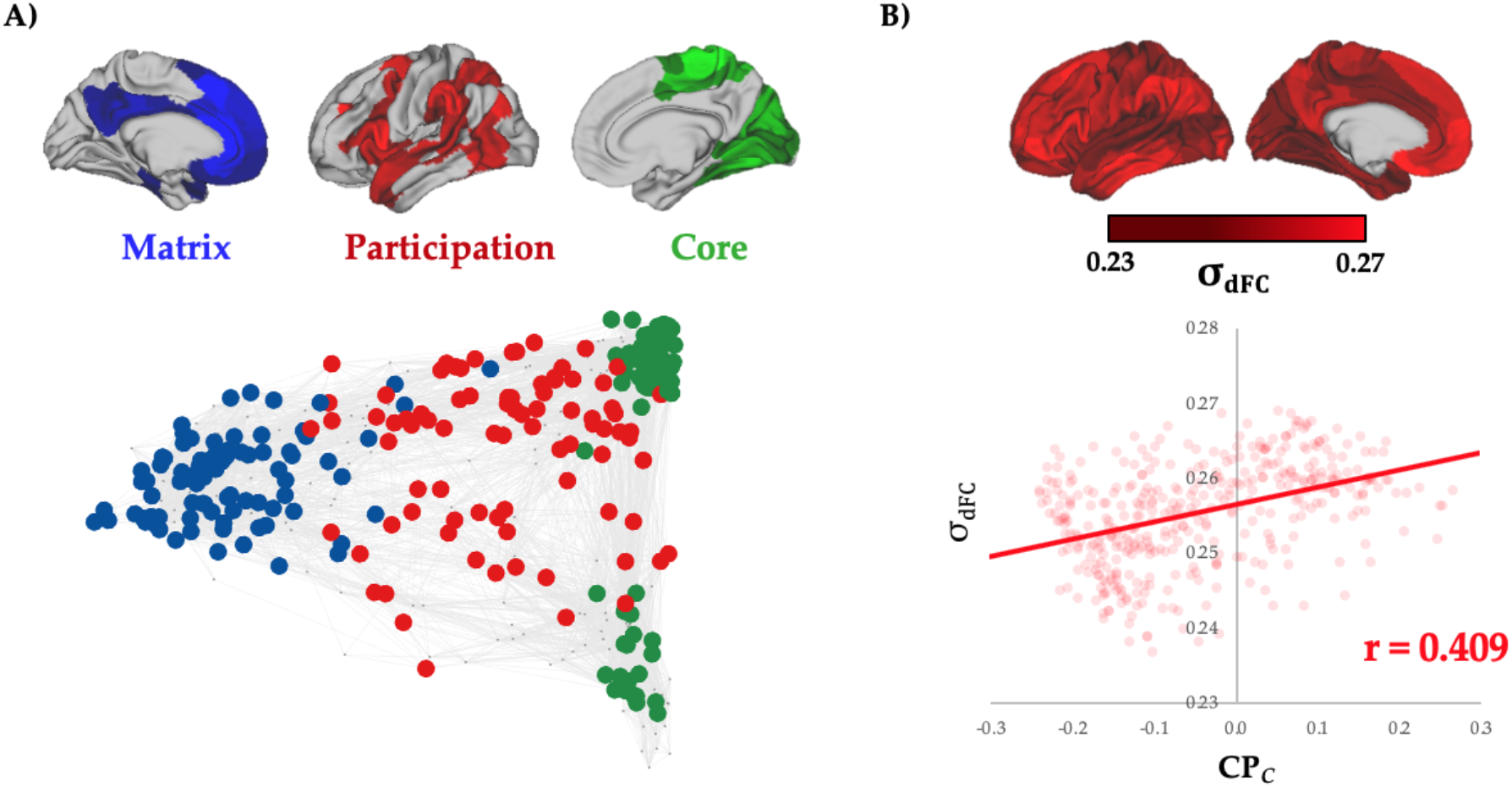
Network Topology. A) Top: parcels within the top 100 ranked regions for each of: Matrix (blue), Participation (red) and Core (green) populations; Bottom: a force-directed plot, in which the top 270 nodes are depicted according to their association with either the Matrix (blue), high Participation regions (red) or Core (green) populations (the remaining 100 regions were plotted as small grey nodes), and edges represent the top 5% of network connections (resting-state functional connectivity); B) scatter plot between the standard deviation of dynamic function connectivity (σ_dFC_) and CP_*C*_ (*r* = 0.409, p_*RAW*_ = 1.6 x 10^-17^; p_*SPIN*_ = 1.0×10^-3^).

Based on the observation that Matrix thalamic nuclei and associative cerebral cortical regions were associated with fluctuations at longer time scales than Core thalamic or sensory cortical regions (i.e., regions with high CP_*T*_/CP_*C*_ had higher *H* and *v.v*.), we predicted that regions with high CP_*C*_ should be associated with more variable connectivity dynamics. In essence, the ability to hold information over longer delays should afford more opportunities for fostering system-level variability. To test this hypothesis, we calculated the variability of windowed time-varying connectivity in the cerebral cortex, and then related this variability to the CP_*C*_ vector. As predicted, we observed a significant, positive correlation between connectivity variability and CP_*C*_ (r = 0.409; p_*RAW*_ = 1.6 x 10^-17^; p_*SPIN*_ = 1.0 x 10^-3^; Figure 5B), suggesting that Matrix-supported cortical regions were associated with more variable connectivity dynamics than the Core-supported cortical regions that are more tethered to sensory inputs (Buckner and Krienen, 2015). This measure was also positively correlated with the first gradient (r = 0.583, p_*RAW*_ = 1.0 x 10^-37^; *p_SPIN_* = 1.0×10^-3^) and both timescale measures (τ: r = 0.657, p_*RAW*_ = 5.8 x 10^-49^; p_*SPIN*_ = 1.0×10^-3^; *H*: r = 0.608, p_*RAW*_ = 7.5 x 10^-42^; p_*SPIN*_ = 1.0×10^-3^) providing further evidence that these features all reflect a low-dimensional organizing principle for the thalamocortical system.

### Reproducibility

The major results were replicated in data from the HCP. In this dataset, CP_*C*_ maps were positively correlated (r = 0.445, p_*RAW*_= 1.3 x 10^-17^), and the replicated CP_*C*_ map also correlated with the first diffusion embedding gradient (r = 0.334, p_*RAW*_ = 4.0×10^-10^) and a longer intrinsic timescale (r = 0.315, p_*RAW*_ = 4.2 x 10^-9^), and inversely correlated with the t1w:t2w map (r = –0.251, p_*RAW*_ = 3.5 x 10^-6^). These results suggest that the thalamocortical measures identified in this study were reliable across independent datasets collected using different scanners and imaging protocols.

## Discussion

In this manuscript, we highlighted a meso-scale organizing principle in the thalamus that is related to low-dimensional (Figure 3), temporal (Figure 4), and topological (Figure 5) patterns that exist within the cerebral cortex during the resting state. This relationship between the cortex and thalamus is consistent with a substantial wealth of both empirical (Garrett et al., 2018; Hwang et al., 2017; Olsen et al., 2012; Rikhye et al., 2018; Schmitt et al., 2017; Shine et al., 2019b) and theoretical (Bell and Shine, 2016; Halassa and Sherman, 2019; Jones, 2009; Sherman, 2007) research, and suggests that an appreciation of subcortical–cortical dynamics is crucial to understanding the organization of the human brain and the way it supports cognition and behaviour.

Our results highlight a number of empirical signatures of cerebral cortical dynamics that relate directly to the mesoscale organization of the thalamus. There are a number of anatomical explanations for these patterns. For one, the thalamus has a vastly lower number of cells than the cortex, suggesting that its engagement would likely foster a relatively low-dimensional cortical architecture (Jones, 2001). Secondly, the different input patterns of the Core and Matrix to the cortex, which project to the granular and supragranular regions, would lead to a relatively feed-forward and feed-back mode of processing (García-Cabezas et al., 2019). This pattern can potentially explain the differences observed in the intrinsic timescales (Figure 4), as feedback processing (via infragranular pyramidal cells inputs into supragranular layers of cortex) is known to occur on slower timescales than feed-forward processing (Bastos et al., 2012; Fries, 2005). Interestingly, there is also evidence that low-dimensional gradients in the cerebral cortex are associated with distinct electrophysiological signals (Hunt et al., 2016), suggesting that the patterns observed in this experiment may also exist at faster time scales than can be accurately measured with BOLD data. Finally, the thalamus represents a major synaptic input to the cortex, and thus likely plays a major role in shaping its resting-state network organization (Bell and Shine, 2016; Hwang et al., 2017; Shine et al., 2019b), both in BOLD data (as presented here) but also in electrophysiological data (Gollo, 2019; Watanabe et al., 2019). Together, these lines of reasoning suggest a putative mechanism for how functional dynamics within the cerebral cortex may be shaped and constrained by activity patterns in the thalamus.

The thalamus has been argued to play a crucial role in shaping brain network topology (Bell and Shine, 2016; Hwang et al., 2017; Shine et al., 2019b). By interconnecting the cerebral cortex with key structures in the subcortex and brainstem, the thalamus likely plays an integrative, hub-like role (Bell and Shine, 2016; Hwang et al., 2017; Shine et al., 2019b). However, a closer inspection of thalamic circuitry suggests that the topological role of the thalamus may be qualitatively distinct from the hubs that exist within the cerebral cortex. For instance, although the thalamus is often identified as a hub in topological analyses of functional brain networks, its intrinsic functional activity is highly damped by pervasive GABAergic input (Jones, 2001). As such, an increase in thalamic activity could promote both integration (by interconnecting otherwise distributed regions of the network) and segregation (by only allowing certain channels of activity to be *on-line* at any one point in time). Our results support this notion, as distinct populations of cells within the thalamus were found to interdigitate between cortical regions organized into distinct topological zones (Figure 5). Further work, particularly in the context of different cognitive tasks (Rikhye et al., 2018; Schmitt et al., 2017; Shine et al., 2019b), is undoubtedly required before the relationships between thalamic sub-populations and cortical network topology can be parsed at the functional level.

With these results in mind, an appreciation of the factors that bias activity within the thalamus become of prime importance. One obvious difference between the Core and Matrix populations is the extent to which they are driven by glutamatergic inputs from the sensory apparatuses, with Core neurons receiving a higher proportion of inputs when compared to Matrix neurons (Jones, 2009; 2001). Inputs from prominent subcortical structures are also known to disproportionately contact the Core and Matrix nuclei, with the deep cerebellar nuclei synapsing with the former, and the basal ganglia with the latter (Kuramoto et al., 2009). Another factor that is of great importance is the neurochemical tone imposed by the ascending arousal system (Shine et al., 2016; Shine et al., 2018). It is well known that different neurotransmitters differentially effect cell populations in the thalamus, with some boosting and others silencing activity within the different nuclei (Llinás and Steriade, 2006; McCormick et al., 2015; 1991; Varela, 2014). Future work that maps the relationships between these neuromodulatory chemicals and cortico-thalamic connectivity will help to clarify the relationships described in this study. The fact that the non-linear intersection between the arousal system, the Matrix thalamus and the supragranular layers of the cortex is causally related to conscious brain activity (Redinbaugh et al., 2019; Suzuki and Larkum, 2020) merely acts to refine the importance of this future work for understanding the rules that govern distributed patterns of higher brain function.

## Conclusion

Our results describe relationships between the relative weighting of distinct cell populations in the thalamus and low-dimensional spatial, temporal, and topological gradients in the cerebral cortex. It is important to note that many other elements within the central nervous system are also organized along spatial gradients, including the striatum (Stanley et al., 2019), cerebellum (Guell et al., 2018), colliculi (Cooper and Rakic, 1981) and hippocampus (Przeździk et al., 2019), suggesting that the interaction between the gradient structure across these diverse systems may represent a key organizing principle for the nervous system (Cisek, 2019; Nieuwenhuys, 1999). In addition, our work highlights the value of linking information from the cellular scale with whole-brain data to understand fundamental principles of brain organization.

## Supplementary Figures

**Figure S1.**
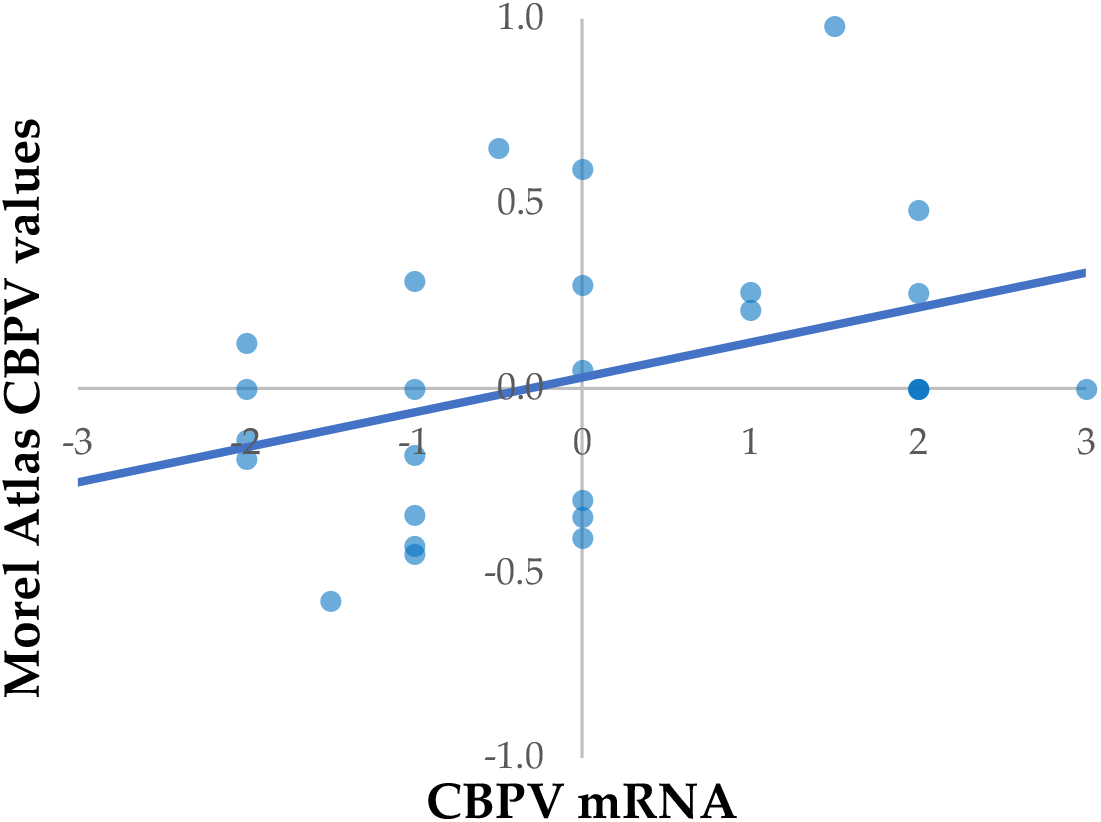
Correlation between Thalamic Calcium Binding Proteins using Allen Human Brain Atlas mRNA and Morel atlas. We observed a positive correspondence between the different measures (r = 0.546; p = 0.003).

**Figure S2.**
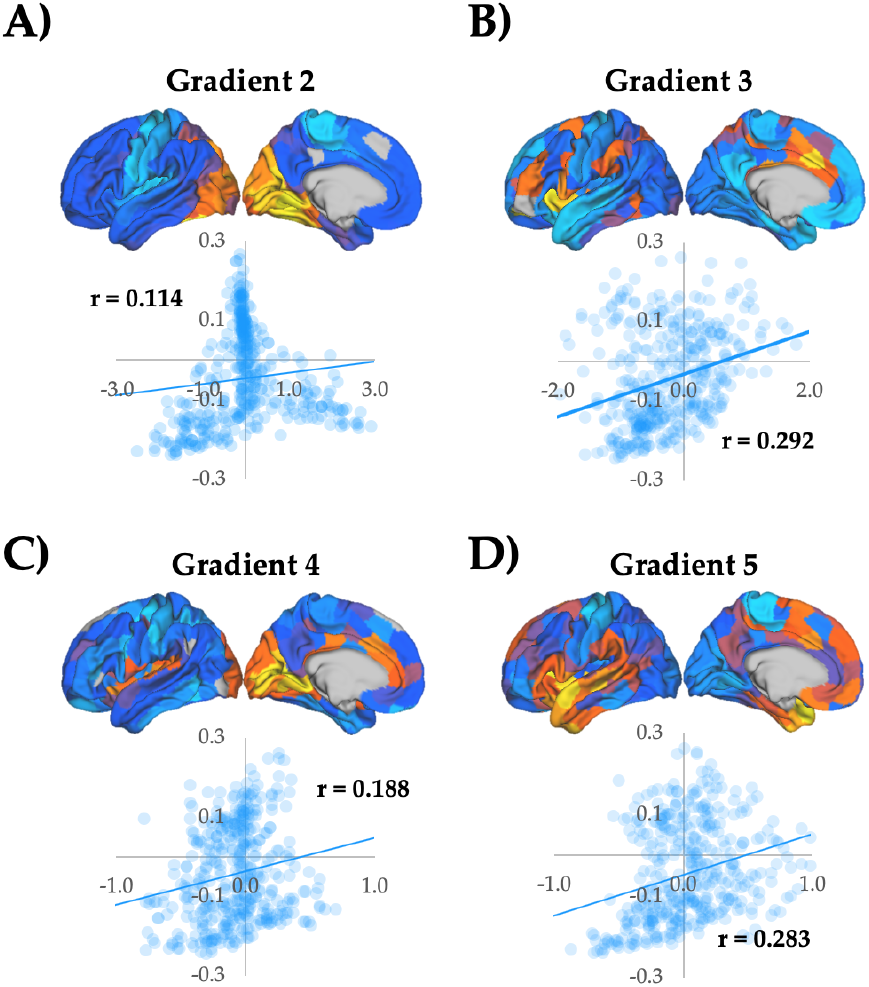
Comparing CP_*C*_ to higher-dimensional gradients. Surface projections and scatter plots relating CP_*C*_ to gradients 2-5 (Margulies et al., 2016). A) Gradient 2: r = 0.114, p_*RAW*_ = 0.02, p_*SPIN*_ = 0.188; B) Gradient 3: r = 0.292, p_*RAW*_ = 2.78 x 10 ^-9^, p_*SPIN*_ = 0.188; C) Gradient 4: r = 0.188, p_*RAW*_ = 2.0 x 10 ^-4^, p_*SPIN*_ = 0.219; D) Gradient 5: r = 0.283, p_*RAW*_ = 8.03 x 10 ^-9^, p_*SPIN*_ = 0.003.

## Acknowledgements

JMS was supported by the University of Sydney Robinson Fellowship and NHMRC GNT1156536.

